# Clonal inactivation of telomerase promotes accelerated stem cell differentiation

**DOI:** 10.1101/2021.04.28.441728

**Authors:** Kazuteru Hasegawa, Yang Zhao, Alina Garbuzov, M. Ryan Corces, Lu Chen, Peggie Cheung, Yuning Wei, Howard Y. Chang, Steven E. Artandi

## Abstract

Telomerase is intimately associated with stem cells and upregulated in cancer, where it serves essential roles through its catalytic action in elongating telomeres, nucleoprotein caps that protect chromosome ends^1^. Overexpression of the telomerase reverse transcriptase (TERT) enhances cell proliferation through telomere-independent means, yet definitive evidence for such a direct role in stem cell function has yet to be revealed through loss-of-function studies. Here, we show that conditional deletion of TERT in spermatogonial stem cells (SSCs) markedly impairs competitive clone formation. Using lineage-tracing from the *Tert* locus, we find that TERT-expressing SSCs yield long-lived clones, but that selective TERT-inactivation in SSCs causes accelerated stem cell differentiation thereby disrupting clone formation. This requirement for TERT in clone formation is bypassed by expression of a catalytically inactive TERT transgene and occurs independently of the canonical telomerase complex. TERT inactivation induces a genome-wide reduction in open chromatin evident in purified SSCs, but not in committed progenitor cells. Loss of TERT causes reduced activity of the MYC oncogene and transgenic expression of MYC in TERT-deleted SSCs efficiently rescues clone formation. These data reveal a required catalytic activity-independent role for TERT in preventing stem cell differentiation, forge a genetic link between TERT and MYC and suggest new means by which TERT may promote tumorigenesis.

## Main Text

Telomerase is enriched in tissue stem cells and activated by somatic promoter mutations in many cancers ^2-4^. The core of the telomerase enzyme is comprised of the catalytic subunit TERT and the telomerase RNA component (*Terc*), a small non-coding RNA scaffold that encodes the template for telomere addition^1^. The critical requirement for telomerase in long-term cell viability is conserved from single cell eukaryotes to humans. Cell proliferation in the absence of telomerase results in a lag phase that is initially well tolerated while telomere reserves are ample. But, proliferation for extended periods in the absence of telomerase culminates in senescence or cell death as telomeres progressively shorten and eventually become dysfunctional. This is particularly evident in laboratory mice, which have very long telomeres (40-80 kb vs 5-15 kb in humans). Telomerase knockout mice are initially viable, but subsequent intergenerational breeding results in severe tissue defects in the advanced generations^5-7^. These findings established the paradigm that TERT is required only for its role in synthesizing telomeres. In contrast with these loss-of-function studies, overexpression studies have sugested that TERT promotes cell proliferation independent of its enzyme function. Conditional transgenic TERT expression caused proliferation of hair follicle stem cells^8^, skin basal layer keratinocytes^9^ and kidney podocytes^10^. These effects of TERT were separable from telomere synthesis because they also occurred with a catalytically inactive TERT allele, or in mice lacking *Terc*^8-10^. In this context, TERT has been shown to activate MYC, WNT and NFkB pathways^8-14^. However, the role of non-canonical functions of TERT in tissue stem cells remains unclear due to the lack of definitive and immediate phenotype in TERT knockout mice.

Tissue homeostasis and carcinogenesis are shaped by cell competition, a mechanism to optimize cell composition in tissues by promoting replacement of damaged or unfit cells with more robust neighboring cells^15^. In renewing mammalian tissues including intestine, skin, and testis, competitive repopulation is a fundamental property of adult stem cells^16^. Stem cell competition results in a dominant expansion of more fit “winner clones” and an elimination of less fit “loser clones” at the niche^17^. This competitive behavior is also characteristic of carcinogenesis during which oncogenic mutations drive cells to clonally expand their territory through a process of super-competition^18^. Among tissues where competitive repopulation has been ovserved, testis shows high telomerase activity and exhibits unusual telomere dynamics in that telomere lengths are preserved with aging, in contrast to the progressive shortening seen in other human tissues. We previously found that spermatogonial stem cells (SSCs) express high levels of TERT and TERT is downregulated with lineage commitment ^7^. To understand the telomere-independent role of TERT in stem cells, we developed a system to mark single SSCs expressing TERT coupled with the ability to conditionally inactivate TERT using a lineage tracing approach. Combining with transgenic rescue experiments and genomic assays, these studies establish TERT as a mediator of stem cell competition in the testis, where it supports stem cell function through a non-canonical mechanism independent of telomere synthesis.

## Results

### High TERT expression marks long-term SSCs that undergo stem cell competition

Spermatogenesis is a dynamic process to produce sperm, composed of mitosis, meiosis and post-meiotic maturation (Extended Data Fig. 1a,b). In the testis, SSCs reside within a functionally and morphologically heterogeneous population undifferentiated spermatogonia (US). Singly isolated A_single_ (A_s_) US undergo incomplete cytokinesis, subsequently producing progressively elongating chains of interconnected cells (A_pr_ - A_16_) (Extended Data Fig. 1b)^19^. Maturation of US yields differentiating spermatogonia (DS), which is accompanied by loss of stem cell potential. To measure *Tert* expression in distinct spermatogonia subpopulations, we purified MCAM^high^ KIT^-^ US (US-h), MCAM^med^ KIT^-^ US (US-m) and MCAM^med^ KIT^+^ DS (Extended Data Fig. 1a-c). Among these subpopulations, *Tert* mRNA expression was high in both US-h and US-m cells, and sharply decreased in DS cells (Extended Data Fig. 1d). *Tert-Tdtomato* reporter mice also showed high Tdtomato expression both in US-h and US-m (Extended Data Fig. 1e). These data indicate that the entire US population exhibits high *Tert* expression.

To functionally study TERT-expressing spermatogonia, we developed a lineage tracing assay using *Tert*^*CreER/+*^ ; *Rosa26*^*lsl-Tdtomato/+*^ (*Tert*^*CreER/+*^) mice, in which TERT-expressing cells are permanently labeled upon tamoxifen-dependent activation of CreER to express Tdtomato (Fig. 1a,b) ^20^. Marking SSCs results in long-lived Tdtomato^+^ clones, also known as patches, comprised of many daughter cells produced by the labeled SSC ^21^. If committed progenitor cells are labeled, only a small transient clone is generated and these cells are lost through differentiation (Fig. 1a,b). Sparse labeling allows rare SSCs to be marked, and in this context each patch derives from a single SSC. Thus, measuring the number of Tdtomato^+^ patches allows a quantitative assessment of stem cell self-renewal activity.

**Fig. 1.**
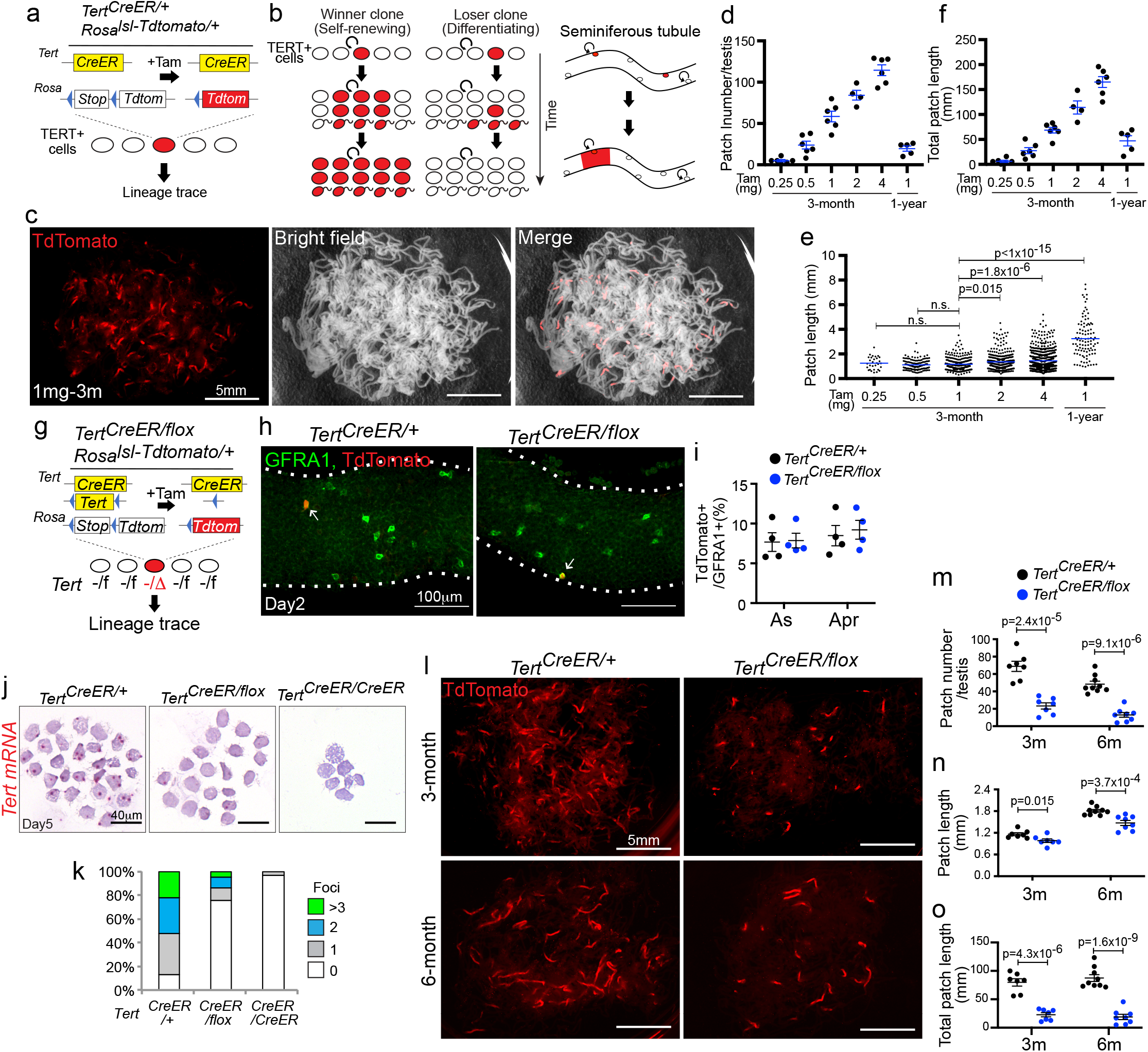
TERT deletion impairs SSC-mediated clone formation. **(a)** *Tert*^*CreER/*+^ *Rosa*^*lsl-Tdtomato/*+^ mice. **(b)** Lineage tracing using *Tert-CreER*. **(c)** Epifluorescence of Tdtomato and bright field image of untangled seminiferous tubules in a single whole testis at 3 months after 1mg tamoxifen injection. Scale bars, 5mm. **(d-f)** Mean patch number (d), mean patch length (e), and total patch length (f) after pulse labeling with indicated tamoxifen dose and span (n=6, 6, 6, 4, 6, 5 mice from left to right). **(g)** *Tert*^*CreER/flox*^*:Rosa*^*lsl-Tdtomato/+*^ mice. **(h)** Whole mount immunofluorescence of GFRA1 and Tdtomato. Scale bars, 100μm. **(i)** Quantification of Tdtomato^+^ cells within GFRA1^+^ A_s_ and A_pr_ cells (n=4 mice per group). **(j)** In situ hybridization against *Tert* mRNA in purified Tdtomato^+^ US at 5 days post-tamoxifen treatment. Undifferentiated spermatogonia from *Tert*^*CreER/CreER*^ mice was used as negative control. Scale bars, 40μm. **(k)** Quantification of foci of *Tert* mRNA (n=232, 243, 341 cells from left to right). **(l)** Epifluorescence of Tdtomato in untangled seminiferous tubules at 3 months and 6 months post-tamoxifen injection. Scale bars, 5mm. **(m-o)** Quantification of mean patch number (m), mean patch length (n), and total patch length (o) (n=7, 7, 9, 8 mice from left to right). Data are represented as Mean ± SEM.

At two days after tamoxifen injection, TdTomato was detected similarly throughout the population of US but not in KIT^+^ cells (Extended Data Fig. 2a-c). At three months, marking TERT^+^ cells yielded labeled patches (Fig. 1c and Extended Data Fig. 2e). As we varied the dose of administered tamoxifen from 0.25 mg to 4 mg, patch number increased in a dose-dependent manner (Fig. 1d, and Extended Data Fig. 2d). Patch length remained constant below 1mg but increased above 2mg, reflecting the fusion of two independently labeled clones with high dose (Fig. 1e). In mice treated with 1mg tamoxifen and traced for one year, patch length increased, clone number decreased, and the total aggregated Tdtomato^+^ patch length (mean patch number times mean patch length) remained constant compared with those traced for three months, consistent with previously described stochastic competition with unlabeled clones (Fig. 1d-f, and Extended Data Fig. 2d) ^21^. At both three months and one year, patches were comprised of cells throughout the spermatogenic lineage (Extended Data Fig. 2f). Altogether, these data show that TERT-expressing SSCs generate long-lived clones and exhibit competition within the stem cell pool.

### TERT deletion impairs SSC-mediated clone formation

To investigate a direct and immediate role for TERT in stem cells, we developed a competitive clone formation assay using *Tert*^*CreER/flox*^*:Rosa*^*lsl-Tdtomato/+*^ (*Tert*^*CreER/flox*^) mice (Fig. 1g, and Extended Data Fig. 3a-c). In this strain, activation of CreER induces Tdtomato labeling and concomitantly inactivates *Tert* in the same TERT^+^ cell, enabling one to trace the fate of cell clones deriving from SSCs in which *Tert* has been somatically deleted in an environment where most neighboring cells retain TERT expression. At two days after 1mg tamoxifen treatment, GFRA1^+^ A_s_ and A_pr_ clones were marked indistinguishably in both *Tert*^*CreER/+*^ and *Tert*^*CreER/flox*^ mice (Fig. 1h,i). Pulse labeling efficiently eliminated *Tert* mRNA in Tdtomato^+^ US (Fig. 1j,k). At three months and six months, deletion of TERT caused a marked reduction in the number of Tdtomato^+^ clones and diminished mean patch length (Fig. 1l-n). Correspondingly, the total aggregated Tdtomato^+^ patch length was sharply reduced (Fig. 1o). Together, these findings indicate that TERT-deletion compromises long-term clone formation ability of SSCs under theses competitive conditions.

### TERT promotes SSC competition independent of its catalytic activity and the telomerase complex

To determine whether the effect of TERT in enhancing SSC competition depends on its catalytic activity, we developed a system to rescue the defect in TERT-deleted cells using tetracycline-regulated TERT transgenes: either one that is wild-type (TetO-TERT) or one that is catalytically inactive (TetO-TERTci) due to a single amino acid substitution in the catalytic site ^9^. We produced compound mouse strains in which activation of CreER simultaneously deletes the floxed *Tert* gene while inducing expression of transgenic *Tert* by triggering expression of the tetracycline transactivator (tTA) that binds and activates the *TetO* promoters (Fig. 2a). We analyzed how TERT loss affected clone formation in *Tert*^*CreER/flox*^*:Rosa*^*lsl-Tdtomato/lsl-tTA*^ (*Tert*^*CreER/flox*^) vs. *Tert*^*CreER/+*^ *:Rosa*^*lsl-Tdtomato/lsl-tTA*^ (*Tert*^*CreER/+*^) controls and we compared these results with restoration of Tert expression in *Tert*^*CreER/flox*^*:Rosa*^*lsl-Tdtomato/lsl-tTA*^:*TetO-Tert* (*Tert*^*CreER/flox*^ *+Tert*) mice or *Tert*^*CreER/flox*^*:Rosa*^*lsl-Tdtomato/lsl-tTA*^:*TetO-Tertci* (*Tert*^*CreER/flox*^ *+Tertci*) mice (Fig. 2a). Transgenic *Tert* and *Tertci* expression was induced at a physiological range by treating mice with low dose doxycycline, which suppresses binding of tTA to the TetO promoter (Extended Data Fig. 4a). Transgenic expression of wild-type *Tert*, but not *Tertci*, restored telomerase activity (Extended Data Fig. 4b). Intriguingly, expression of either *Tert* or *Tertci* transgenes fully rescued patch number, average patch length, and total patch length (Fig. 2b-e). Furthermore, expression of TERTci significantly increased total patch length compared to control *Tert*^*CreER/+*^ mice (Fig. 2e). These results indicate that TERT promotes enhanced stem cell competition independent of its catalytic activity.

**Fig. 2.**
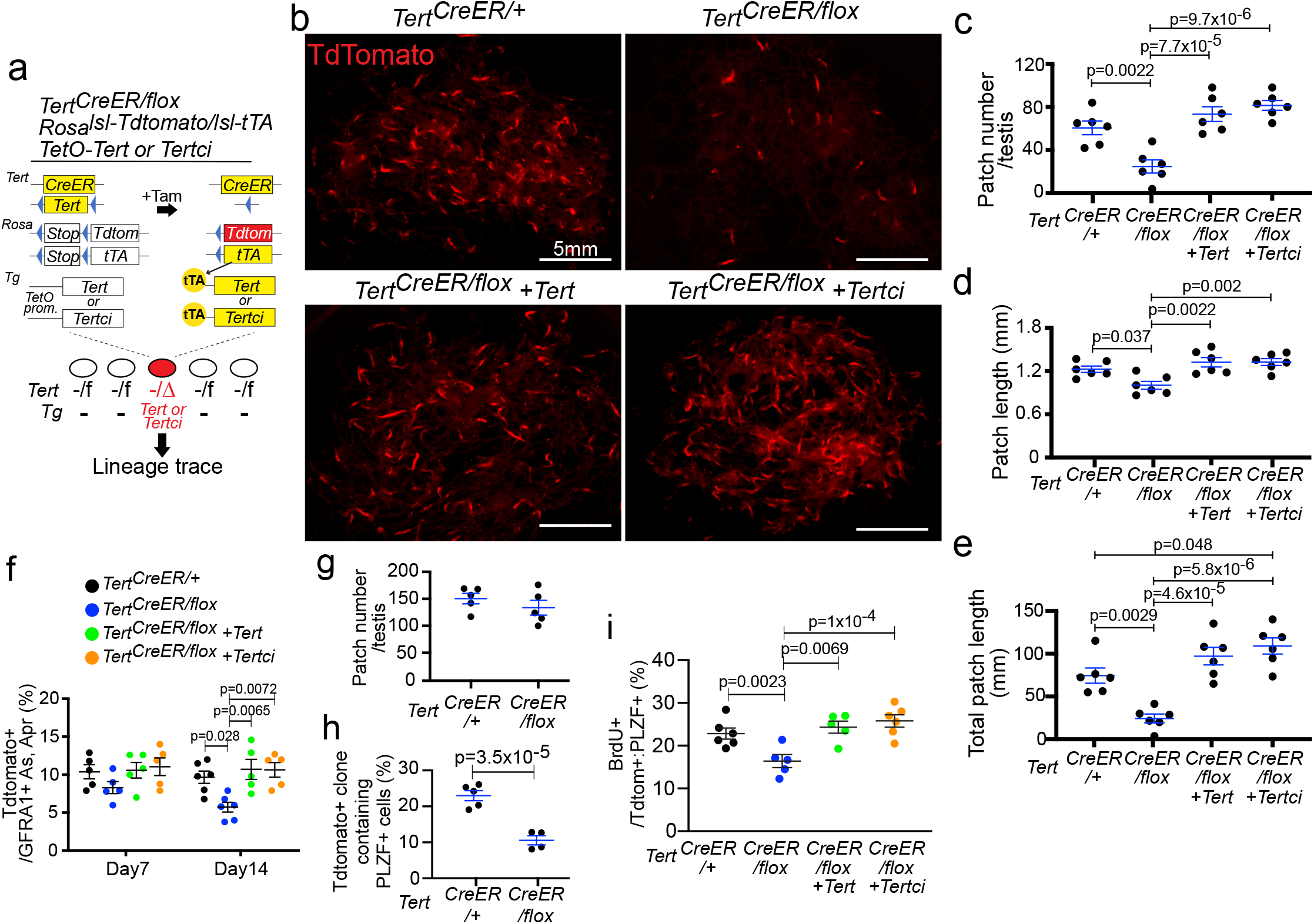
TERT promotes SSC competition independent of its catalytic activity and the telomerase complex. **(a)** *Tert*^*CreER/flox*^*:Rosa*^*lsl-Tdtomato/lsl-tTA:*^ *TetO-Tert or -Tertci* mice. **(b)** Epifluorescence for Tdtomato in untangled seminiferous tubules at 3 months after tamoxifen injection. Scale bars, 5mm. **(c-e)** Quantification of mean patch number (c), mean patch length (d), and total patch length (e) (n=6 mice per group). **(f)** Quantification of TdTomato^+^, GFRA1^+^ A_s_ and A_pr_ cells at 7 or 14 days post labeling (n=5, 5, 5, 5, 6, 6, 5, 5 mice from left to right). **(g)** Mean patch number at 6 weeks after tamoxifen injection (n=5 mice per group). **(h)** Quantification of percent Tdtomato^+^ clones containing PLZF^+^ cells (n=5,4 mice from left to right). **(i)** Quantification of BrdU^+^ cells in Tdtomato^+^, PLZF^+^ undifferentiated spermatogonia (n=6, 5, 5, 6 mice from left to right). Data are represented as Mean ± SEM.

*Terc* encodes the RNA template for telomere addition and serves as the central scaffold for assembly of the telomerase complex. Therefore in the absence of *Terc*, TERT and the other components of telomerase fail to associate ^1,22^. To determine whether formation of the telomerase complex is required for the TERT-dependent SSC clone formation, we produced first generation *Terc*^-/-^:*Tert*^*CreER/flox*^ *:Rosa*^*lsl-Tdtomato/+*^ (G1 *Terc*^-/-^:*Tert*^*CreER/flox*^) mice (Extended Data Fig. 4c. In this strain, telomerase was inactive in testes (Extended Data Fig. 4d). Three months after tamoxifen treatment, conditional deletion of *Tert* in *Terc*-deficient mice impaired patch number, patch length, and total patch length (Extended Data Fig. 4e -h). These findings uncouple the requirement for TERT in stem cell competition from engagement in the classical telomerase complex.

Somatic deletion of *Tert* in mice with very long telomeres is unlikely to cause telomere dysfunction ^7,23,24^. When telomeres become short, however, a DNA damage response activates the p53 tumor suppressor protein, triggering either cell death or senescence ^25^. To understand whether the impaired clone formation in TERT-deleted SSCs is mediated by p53, we generated *Trp53*^*flox/flox*^*:Tert*^*CreER/flox*^*:Rosa*^*lsl-Tdtomato/+*^ mice and examined competitive clone formation (Extended Data Fig. 5a). At day five, the deletion efficiency of the *Trp53-flox* alleles in Tdtomato^+^ US was 44.4% (Extended Data Fig. 5b,c). We found that deletion of p53 failed to rescue the impaired clone formation associated with *Tert* inactivation, as measured by patch number, patch length, and total patch length (Extended Data Fig. 5d-g). Consistent with these findings, we found no accumulation of γH2AX, a marker of DNA damage, in *Tert*^*CreER/flox*^ mice, whereas γH2AX was elevated in G6 *Tert*^*Tdtomato/Tdtomato*^ mice with dysfunctional telomeres (Extended Data Fig. 5h,i). Taken together, the effects of TERT in promoting stem cell-derived clone formation is independent of catalytic activity, the DNA damage response and the canonical telomerase complex.

### Accelerated differentiation and impaired proliferation of SSCs lacking TERT

The preferential elimination of conditionally TERT-deleted SSCs could be caused by accelerated differentiation, impaired proliferation, or increased apoptosis. To distinguish among these possibilities, we investigated how clonal deletion of TERT influences SSC fate across at time points. At 14 days after tamoxifen administration, Tdtomato^+^ A_s_ and A_pr_ US were significantly decreased in *Tert*^*CreER/flox*^ mice, and the reduction of those cells was rescued by transgenic expression of either TERT or TERTci (Fig. 2f and Extended Data Fig. 6a). At six weeks when a comparable frequency of labeled patches were found in *Tert*^*CreER/+*^ and *Tert*^*CreER/flox*^ mice, labeled clones harbored significantly fewer US (Fig. 2g,h Extended Data Fig. 6b,c). These results indicate that TERT deletion promotes differentiation of US. To understand how TERT loss affects cell proliferation, we measured BrdU incorporation seven days after tamoxifen administration. The percentage of US incorporating BrdU was significantly reduced in *Tert*^*CreER/flox*^ animals and proliferation was rescued with transgenic expression of *Tert* or *Tertci* (Fig. 2i and Extended Data Fig. 6d). There was no increase in apoptosis by cleaved-PARP staining in *Tert*^*CreER/flox*^ mice, but apoptosis was elevated in testes from G6 *Tert*^*Tdtomato/Tdtomato*^ mice with critically short telomeres (Extended Data Fig. 6e,f). Taken together, impaired clone formation of TERT deleted SSCs is caused by accelerated differentiation and decreased proliferation.

### TERT deletion compromises chromatin accessibility in SSCs but not in committed progenitors

To understand how TERT deletion affects global chromatin structure, we performed ATAC-seq, which allows an assessment of chromatin accessibility genome-wide ^26^. To first define the patterns of chromatin changes during normal spermatogenesis, we purified US-h, US-m, and DS from *Tert*^*CreER/+*^ mice that had been injected with tamoxifen seven days prior to isolation, while spermatocytes (SP) and round spermatids (RS) were purified based on differential *Tert* promoter activity in *Tert*^*Tdtomato/+*^ mice ^7^. Principal component analysis (PCA) revealed that US-h and US-m clustered together in the lower left quadrant, consistent with their similar gene expression patterns (Extended Data Fig. 7a) ^27^. DS cells localized in the upper left quadrant, while SP and RS populations were clustered together in the right lower quadrant (Extended Data Fig. 7a), indicating that the PC1 axis captures the changes in global chromatin state associated with differentiation. Similarly, Pearson correlation hierarchical clustering showed a high correlation in open chromatin patterns among spermatogonia subpopulations, but abrupt changes of chromatin accessibility globally upon entry to meiosis (Extended Data Fig. 7b). The number of unique ATAC-seq peaks and promoter chromatin accessibility around transcription start sites were highest in US-h and US-m and decreased significantly during differentiation into DS and SP (Fig. 3a,b, and Extended Data Fig. 7c,d). The promoter region of *Tert* was accessible in US-h, US-m and DS, but inaccessible in SP and RS, consistent with the expression pattern of TERT during spermatogenesis (Extended Data Fig. 8a) ^7^. Altogether, these data reveal that SSCs show a marked increase in chromatin accessibility, indicating that a global reduction in chromatin accessibility occurs during spermatogenesis.

**Fig. 3.**
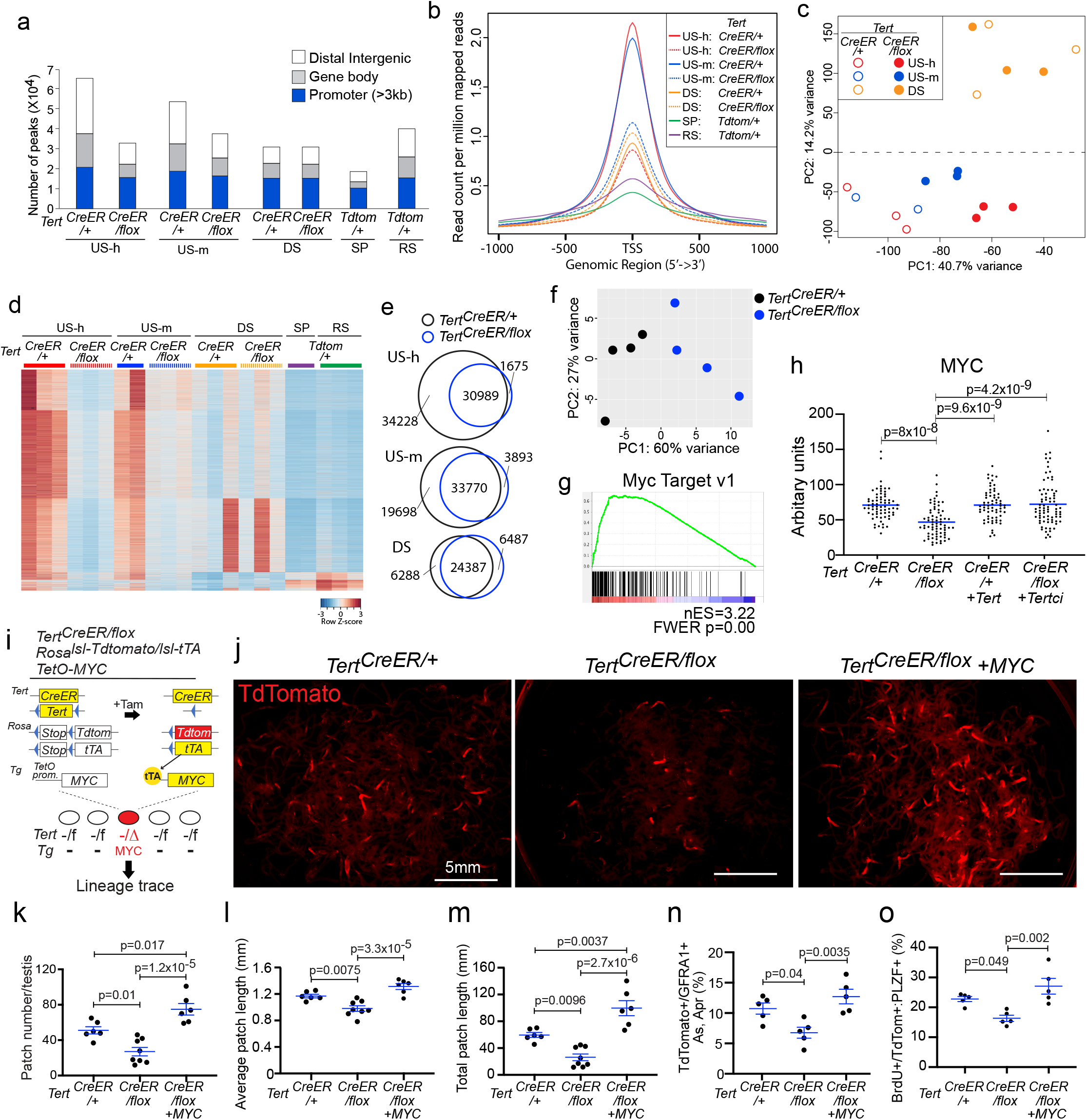
TERT deletion compromises chromatin accessibility in SSCs. **(a)** Peak calls from ATAC-seq data. Peak calls from each cell type are shown individually. Color indicates the type of genomic region overlapped by the peak. **(b)** Average tag density of ATAC-seq reads around transcription start sites. **(c)** PCA of ATAC-seq data from US-h, US-m and DS cells purified from *Tert*^*CreER/+*^ versus *Tert*^*CreER/flox*^ mice. **(d)** Heatmap representation of 11656 peaks that are significantly different between *Tert*^*CreER/+*^ and *Tert*^*CreER/flox*^ in US-h, US-m, and DS or significantly more open either in SP or RS. Each row represents one ATAC-seq peak. Color represents the relative ATAC-seq accessibility. **(e)** Venn diagrams of the peaks in US-h, US-m and DS cells from *Tert*^*CreER/+*^ and *Tert*^*CreER/flox*^ mice.

To understand how conditional TERT-deletion influences chromatin accessibility in SSC populations, we performed ATAC-seq on US-h, US-m and DS isolated from *Tert*^*CreER/flox*^ mice. Pearson correlation hierarchical clustering showed that the overall pattern of chromatin accessibility in TERT-deleted cells remained similar to that of *Tert*^*CreER/+*^ controls (Extended Data Fig. 7b). However, PCA revealed that TERT deletion caused a shift in the US-h and US-m along the differentiation axis, whereas TERT loss had no discernible effect on committed DS (Fig. 3c). Deletion of TERT in the US populations caused a marked reduction in the number of open chromatin peaks and diminished chromatin accessibility surrounding TSSs to a level resembling the control DS (Fig. 3a,b,d,e and Extended Data Fig. 7e). In contrast, peak number was unaffected by TERT deletion in DS (Fig. 3 a,b,d,e and Extended Data Fig. 7e). This reduction in open chromatin peaks was evident in genes associated with stemness in SSCs, including *Ret, Gfra1, Cdh1*, and *Zbtb16*, whereas those associated with differentiation including *Kit, Prm2*, and *Prm3* remained unchanged (Extended Data Fig. 8b-d). Pathway analysis revealed that genes in the MAPK signaling pathway, which promotes self-renewal of SSCs ^28^, were particularly enriched among those showing loss of open chromatin peaks in the TERT-deleted undifferentiated spermatogonia (Extended Data Fig. 7f). Taken together, conditional inactivation of TERT caused a loss of open chromatin selectively in the stem cell containing population but not in committed progenitors, consistent with accelerated stem cell differentiation caused by TERT deletion.

### TERT promotes competitive clone formation of SSCs through MYC

To understand how TERT promotes competitive clone formation in SSCs, we examined gene expression in TERT-deleted US-h cells by RNA-seq seven days after tamoxifen administration. PCA showed that TERT-deleted US-h clustered separately from TERT^+^ controls (Fig. 4a). Using strict cutoffs for significance, there were 23 genes downregulated and 116 genes upregulated in TERT-deleted US-h (Extended Data Fig. 9a). Gene set enrichment analysis revealed that spermatogenesis-related genes were upregulated in TERT-deleted US-h, consistent with their accelerated differentiation (Extended Data Fig. 9b). Several gene sets were downregulated in TERT-deleted US-h cells, including E2F targets and G2M checkpoints, reflecting the quantitative reduction in proliferation in these cells (Extended Data Fig. 9c). The most significantly downregulated gene set was ‘MYC targets v1’ and a second gene set ‘MYC targets v2’ was also represented (Fig. 3b, and Extended Data Fig. 9c). Consistent with this, MYC protein was significantly decreased in TERT-deleted US and MYC levels were restored by transgenic expression of TERT or TERTci (Fig.4c, and Extended Data Fig.9d,e). *Myc* mRNA levels remained unchanged, suggesting that TERT promotes MYC expression at the post-transcriptional level (Extended Data Fig. 9f).

**Fig. 4.**
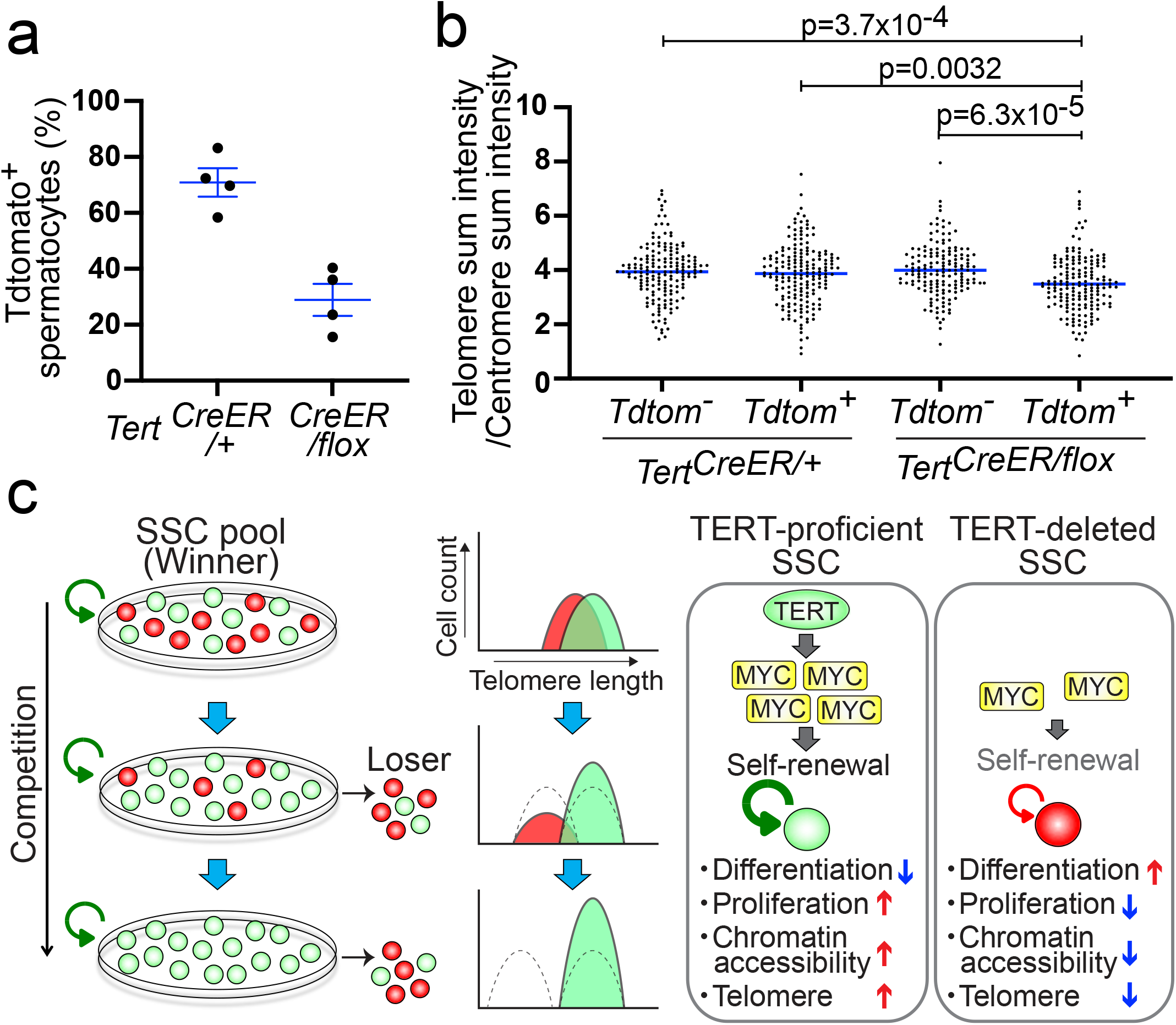
MYC rescues competitive clone formation of TERT-deleted SSCs. **(a)** PCA of RNA-seq data from Tdtomato^+^ US-h. **(b)** Significant down-regulation of MYC target genes in *Tert*^*CreER/flox*^ US-h cells. RNA-seq data was analyzed using gene set enrichment analysis. **(c)** Quantification of mean signal intensity for MYC staining in PLZF^+^ undifferentiated spermatogonia using testicular cross sections at 7 days after labeling. **(d)** *Tert*^*CreER/flox*^*:Rosa*^*lsl-Tdtomato/lsl-tTA*^: *TetO-hMYC* mice. **(e)** Epifluorescence for Tdtomato in untangled seminiferous tubules. Scale bars, 5mm. **(f to h)** Quantification of mean patch number (f), mean patch length (g), and total patch length (h) (n=6, 8, 6 mice from left to right). **(i)** Quantification of Tdtomato^+^ GFRA1^+^ A_s_ and A_pr_ clones at 14 days post labeling detected by whole mount immunofluorescence (n=5 mice per group). **(j)** Quantitative analysis of Tdtomato^+^ BrdU^+^ PLZF^+^ undifferentiated spermatogonia using testicular cross sections at 7 days post labeling (n=5 mice per gropu). Data are represented as Mean ± SEM.

MYC is a transcription factor that promotes cell competitiveness by regulating cell proliferation, growth, and metabolism ^29-32^. MYC has been shown to promote SSC self-renewal and also an important oncogene ^33-35^. Given the genetic links between TERT and MYC, we hypothesized that MYC overexpression might rescue the failure of clone formation in TERT-deleted SSCs. To test this idea, we intercrossed *Tert*^*CreER/flox*^: *Rosa*^*lsl-Tdtomato/lsl-tTA*^ mice with *TetO-human MYC* transgenic mice (*Tert*^*CreER/flox*^ *+MYC*) (Fig. 4d). This system allows simultaneous deletion of the residual TERT allele and activation of transgenic MYC selectively in a lineage of TERT-expressing stem cells. To limit expression levels of transgenic MYC, mice were treated with doxycycline, which reduced transgenic *MYC* mRNA levels by 5.6 fold (Extended Data Fig. 9g). Three months after tamoxifen treatment, the defects in clone formation associated with TERT loss were dramatically rescued by MYC expression, as measured by patch number, patch length and total patch length (Fig. 4e-h). MYC expression also restored the number of GFRA1^+^ cells, proliferation and expression of MYC target genes (Fig. 4i,j and Extended Data Fig. 9h). Taken together, these results indicate that the marked defect in stem cell competition associated with TERT loss is due to impaired MYC function and establish an epistatic relationship between TERT and MYC.

## Discussion

TERT is expressed in many stem cell compartments governed by competitive repopulations and TERT upregulation is selected for during human carcinogenesis. By tracing the fate of individual SSCs *in vivo*, we found that loss of TERT severely compromises competitive clone formation and accelerates stem cell differentiation. In this context, TERT-deleted stem cells represent “loser” clones outcompeted by TERT-proficient “winner” clones. This requirement for TERT in competitive clone formation is independent of telomeres and TERT catalytic function; instead, TERT maintains chromatin accesiblity across many genes expressed in stem cells. These findings establish TERT as a key determinant regulating competition between tissue stem cells by favoring self-renewal and disfavoring differentiation (Extended Data Fig. 10). This result is surprising in that germline inactivation of TERT in mice is well tolerated initially, although SSC function is ultimately compromised by telomere dysfunction after many generations of telomerase-deficiency. Competitive behavior of SSCs may expain why germline mutations in *Tert* do not cause an obvious phenotype in laboratory mice. Selection of winner clones and elimination of loser clones in the testis may be determined in part based on competition for limiting niche signals ^36^. This niche-dependent competition between TERT-proficient and TERT-deficient cells may amplify cell-autonomous defects caused by acute TERT deletion, while promoting elimination of TERT-deleted SSCs. Homogeneous deletion of TERT throughout the organism does not create a TERT-dependent competition, and in this context TERT-deficient cells are capable of self-renewal and differentiation to produce sperm. Thus, germline gene inactivation identifies only one layer of activity, whereas clonal, conditional gene deletion can reveal deeper aspects of gene function.

While TERT levels are important in maintaining stem cell function, they may play a similar role during carcinogenesis. TERT is upregulated in many human tumors through non-coding mutations in the proximal TERT promoter. These TERT promoter mutations show strong positive selection during tumor progression, as the prevalence of these mutations increases substantially from pre-invasive to invasive stages. Our data establishing an epistatic relationship between TERT and MYC provide support for this model. MYC activity is central in determining outcomes in cell competition; cells expressing higher MYC outcompete those with lower MYC and MYC represents a key node in many human cancers ^29-32^. TERT upregulation in cancer likely increases telomerase activity and stabilizes telomeres, thereby preventing the negative effects of dysfunctional telomeres on cell cycle and cell survival. At the same time, TERT upregulation may enhance cell competition through MYC and its non-canonical effects in promoting self-renewal and preventing differentiation. MYC can also control TERT levels by directly activating transcription of *Tert* through promoter binding ^37-39^. Therefore TERT and MYC may comprise a positive feedback loop that promotes cell competition in various tissues during development, homeostasis and carcinogenesis. Our findings using clonal deletion in SSCs add important new complexity to our understanding of TERT function. Taken together, this study establish a role for TERT in promoting clonal competition in stem cells *in vivo* with important implications for understanding tissue homeostasis, cancer development and telomere maintenance.

## Methods

### Animals

All animal protocols were approved by the Institutional Animal Care and Use Committee at Stanford University. All experiments are in compliance with the ethical regulations of Stanford University. *Tert-CreER, TetO-Tert, TetO-Tertci, Tert-Tdtomato* mice were previously reported ^7,10,20^. *Rosa-lslTdtomato* ^40^, *Rosa-lsl-tTA* ^41^, *TetO-hMYC* ^42^, *Terc-KO* ^43^, *Trp53-flox* ^44^ mice were purchased from The Jackson laboratory. Tamoxifen (Cayman) was dissolved in corn oil (Sigma-Aldrich) at 5 - 20mg/ml by incubating at 50 °C for 30min with mixing every 5min. Two- to three-month-old mice were administrated with 0.25 - 4mg per 25g body weight tamoxifen by oral gavage or intra-peritoneal injection. Doxycycline (Sigma-Aldrich) was dissolved in drinking water in light-protected bottles at 1 or 3 μg/ml and changed every 4 days. BrdU (Sigma-Aldrich) was dissolved in PBS at 10mg/ml and intra-peritoneally injected at 1.25mg per 25g body weight at 2 hours before sacrifice.

### Generation of TERT-flox mice

9kb fragment of TERT locus was subcloned and Lox-Puro-lox cassette from pBS.DAT-LoxStop plasmid (kindly gifted from David Tuveson) was inserted at the BsiWI site in the second intron. Another loxP sequence and NdeI site were inserted at the KasI site in the sixth intron. The targeting vector was linearized and electroporated into J1 mouse ES cells. After positive selection with puromycin, correctly targeted ES clones were selected by long-range PCR and Southern blotting, and then injected into C57BL6 blastocysts to generate the knock-in line. To remove floxed Puro cassette, the knock-in line was crossed with CMV-cre mice ^45^ and puro-negative TERT-floxed mice were selected by PCR and Southern blotting using genomic DNA from tail tips. *TERTflox/+* mice were born at normal Mendelian frequency.

### Lineage tracing assay

After tamoxifen injection, testes were detunicated, dissociated using fine forceps in PBS containing 1mg/ml collagenase IV (Worthington) for 10min to remove interstitial cells, and placed in cold PBS. Images were captured with a fluorescent dissection microscope and the patch number and length were measured with ImageJ. Total patch length was calculated by multiplying the patch number by the average patch length.

### Whole-mount Immunofluorescence of seminiferous Tubules

Seminiferous tubules were dissociated using fine forceps in PBS containing 1mg/ml collagenase IV for 10min, fixed with 4%PFA at 4°C for 2h, cleared with 0.1% Igepal CA-630 (Sigma-Aldrich) in PBST, and dehydrated and rehydrated by immersing in a gradient of methanol diluted with PBST (25%, 50%, 75%, 100%, 75%, 50%, 25%) at 4°C for 5min each. After washing in PBST, tubules were incubated in blocking buffer (0.5% BSA PBST) (Sigma-Aldrich), followed by incubation with antibodies in Immuno shot immunostaining Mild (Cosmo Bio) at 4°C for two days. After extensive wash with PBST, tubules were incubated with secondary antibodies in blocking buffer at room temperature for 90min, washed with PBST, and then mounted in Vectashield with DAPI (Vector laboratories). Images were captured on a Leica SP5 confocal microscope and processed in Photoshop.

### Section immunostaining

Testes were detunicated, fixed with 4% PFA at 4°C overnight, incubated in a gradient ethanol, xylen, embedded in paraffin, and cut into 5-μm sections. After rehydration, antigen retrieval using Antigen retrieval solution citric acid or tris-based (Vector) for 10min in a pressure cooker. Sections were blocked, incubated with primary antibody at 4°C overnight. After washing with PBS, sections were incubated with secondary antibodies at room temperature for 1h and mounted in Vectashield with DAPI. For co-staining using rabbit anti-RFP antibodies and rabbit anti-PLZF antibodies, sections were antigen retrieved, incubated with anti-RFP antibody, then with HRP-conjugated anti-rabbit secondary antibody as described above, and signals were detected with TSA plus Cyanine 3 system (Perkin Elmer). Those antibodies were stripped off by antigen retrieval, and sections were further stained with BrdU, cleaved-PARP, or PLZF antibodies. For BrdU detection, those slides were treated with 2N HCl for 20min, incubated with rabbit ant-PLZF and rat anti-BrdU antibodies at 4°C overnight, and signals were detected by Alexa488-conjugated anti-rat IgG and Cy5-conjugated anti-rabbit IgG antibodies. For cleaved-PARP detection, slides were incubated with rabbit ant-PLZF and mouse anti-cleaved PARP antibodies at 4°C overnight, and signals were detected by Alexa488-conjugated anti-mouse IgG and Cy5-conjugated anti-rabbit IgG antibodies. For triple staining using rabbit anti-RFP, rabbit anti-PLZF, and rabbit c-MYC antibodies, sections were stained with anti-Tdtomato antibody using TSA plus Cyanine 3 system, and antibodies were stripped off. Then those sections were stained with anti-MYC antibody with TSA plus fluorescein system (Perkin Elmer) followed by antigen retrieval to remove antibodies. Finally, the sections were further stained with anti-PLZF antibody and Cy5 conjugated anti-rabbit IgG. Slides were mounted in Vectashield with DAPI. Images were captured on a fluorescent microscope and processed in Photoshop. Signal intensity of c-MYC signal was quantified with Image J.

### TRAP assays (Telomeric Repeat Amplification Protocol)

A two-step TRAP procedure was performed as previously reported^46^. Extract fractions from whole testis at 3 weeks or FACS-sorted undifferentiated spermatogonia were incubated with telomeric primers for a 30 min initial extension step at 30°C in a PCR machine, followed by 5 min inactivation at 72°C. Without purification, 1μl of the extended reaction was PCR amplified (cycles of 30 seconds at 94°C, followed by 30 seconds at 59°C) in presence of ^32^P end-labeled telomeric primers that has been purified using a micro-spin G-25 column (GE healthcare). PCR reactions were resolved by 9% polyacrylamide gel electrophoresis at room temperature, and the gel was exposed to a phosphor-imager and scanned by a Typhoon scanner. The scanned image was quantitated using the TotalLab Quant software. Representative gel images were presented among at least 2 repeats.

### FACS analysis

Testes were detunicated, lightly dissociated in PBS, and incubated in PBS containing DNase I (Worthington) and 1mg/ml collagenase I (Worthington) at 32°C for 10 min. Cells were centrifuged at 250g for 5min and supernatant was removed. After repeating collagenase I treatment, testicular cells were further digested with TrypLE Express (GIBCO) at 32°C for 15 min. During enzymatic digestions, seminiferous tubules were mechanically fragmented with vigorous pipetting every 5min. Cells were sequentially filtered with 70 μm and 40 μm strainers, resuspended in cold FACS buffer (2% FBS, 1mM EDTA in PBS), and incubated with antibodies on ice for 30min. After PBS wash, cells were resuspended in cold FACS buffer containing DAPI, and analyzed and sorted with a BD Aria II (BD Biosciences). Data was analyzed with FlowJo software. The list of antibodies is available in supplemental table.

### RNA in situ hybridization

5 days post labeling, testes were collected and Tdtomato+ undifferentiated spermatogonia were FACS-sorted, and cytospun at 300 r.p.m. for 5 min onto slides. Slides were fixed in 4% (v/v) PFA for 30 min at room temperature and processed for single-molecule RNA FISH using RNAscope 2.5 HD Reagent KIT-RED (Advanced Cell Diagnostics) and probes against mouse *Tert* or *Trp53* (Advanced Cell Diagnostics) according to the manufacturer’s instructions.

### Quantitative RT-PCR

For qRT-PCR, cells were directly sorted into Trizol LS (Thermo Fisher Scientific) by FACS and mixed with 100ng yeast tRNA as carrier. RNA was purified Direct-zol RNA Microprep (Zymo Research) and cDNA was synthesized using oligo-dT and SuperScript IV First-Strand Synthesis system (Thermo Fisher Scientific). For qRT-PCR of *Tert* and mouse *Myc*, Universal Probe Library Probe #066 and #72 (Roche) were used, respectively. For these reactions, TaqMan Fast Advanced Master Mix (Thermo Fisher Scientific) were used. For other qRT-PCR, PowerUp SYB Green Master Mix (Thermo Fisher Scientific) was used according to the product manual. PCR analysis was done with a 7900HT Fast Real-Time PCR System machine (ABI). List of primers is available in supplemental table.

### RNA-sequencing

US-h cells were directly sorted into Trizol LS by FACS and RNA was purified using Direct-zol RNA Microprep. Genomic DNA was digested with on-column DNase treatment. RNA quality was checked by Bioanalyzer 2100 (Agilent). RNA-seq libraries were constructed using SMARTer Stranded Total RNA-seq Kit v2 – Pico Input Mammalian (Clontech), starting from 5ng total RNA. cDNA was synthesized and amplified according to the manual. After the rRNA removal step, cDNA was amplified with 13 cycles of PCR reactions. Quality of purified cDNA libraries was confirmed by Bioanalyzer 2100 and concentration of cDNA. Libraries were sequenced on the Illumina NextSeq platform, generating about 16-24 million 75-bp paired-end reads per library. Four biological replicates per sample were analyzed. Raw reads were trimmed by TrimGalore 0.4.0 (Babraham Bioinformatics), mapped to mm10 by tophat 2.0.13 ^47^, analyzed by the DEseq2 packages ^48^.

### ATAC-sequencing

ATAC-seq libraries were made as described previously ^49^ using the Omni-ATAC protocol. Adjustments to the protocol were made to reflect two primary features of the cell types profiled in this work. First, the amount of Tn5 transposase added to each reaction was modulated to maintain proportionality with the number of cells assayed. For example, a normal reaction uses 50,000 cells and 2.5 ul of Tn5 transposase in a 50 ul reaction. In the case of rarer spermatogonial stem cells, only 5,000 cells could be obtained so only 0.25 ul of Tn5 transposase was used in a 50 ul reaction. The difference in volume was adjusted using water. Second, the ploidy of each cell type was taken into account and the amount of Tn5 was adjusted based on ploidy as well. For example, round spermatid cells are haploid, so transposition of 50,000 cells would require 1.25 ul of Tn5 transposase in a 50 ul reaction. Similarly, spermatocytes are 4N meiotic cells so the amount of Tn5 transposase was increased proportionately and the amount of water in the reaction was reduced. In all cases, regardless of cell number or ploidy, the reaction volume of the transposition reaction was kept constant at 50 ul. All ATAC-seq reactions were performed using homemade Tn5 transposase and Tagment DNA (TD) buffer ^50^. Downstream amplification and purification of libraries was performed as described previously ^26,49,51^.

ATAC-seq data pre-processing was completed using the PEPATAC pipeline (http://code.databio.org/PEPATAC/). The mm10 genome build (https://github.com/databio/refgenie) was used for alignment. Briefly, all fastq files were first trimmed to remove Illumina Nextera adapter sequence using Skewer ^52^ with “-f sanger –t 20 –m pe –x” options. FastQC (http://www.bioinformatics.bbsrc.ac.uk/projects/fastqc) was used to validate proper trimming and check overall sequence data quality. Bowtie2 ^53^ was then used for pre-alignments to remove reads that would map to chrM (revised Cambridge Reference Sequence), alpha satellite repeats, Alu repeats, ribosomal DNA repeats, and other repeat regions with “-k 1 -D 20 -R 3 -N 1 -L 20 -i S,1,0.50 -X 2000 –rg-id” options. Bowtie2 was then used to align to the mm10 reference genome using “--very-sensitive -X 2000 --rg-id” options. Samtools ^54^ was used to sort and isolate uniquely mapped reads using “-f 2 -q 10 -b -@ 20” options. Picard (http://broadinstitute.github.io/picard/) was used to remove duplicates. Then the bam files were merged by conditions, and MAC2 ^55^ was used to call peaks with parameter “-q 0.05 --nomodel--shift 0”. The narrow peaks were then filtered by the ENCODE 7 hg19 blacklist, as well as peaks that extend beyond the ends of chromosomes. Bedtools ^56^ was used to retrieve the reads of the called peaks for each sample with multicov module. All the samples have similar sequencing depth, mitochondrial rate, and duplication rate. The spermatocyte and the round spermatid samples have similar sequencing depth compared to all other samples, but slightly higher mitochondrial rate and lower duplication rate, so have more final reads after initial processing and filtering. To make all the samples comparable for the statistical analysis, we used final reads as the normalization factor. R package “DESeq2” ^48^ was used for statistical analysis to identify significant peaks between different conditions. The differential peaks were called between US-h CreER/+ and US-h CreER/flox samples. Peaks with FDR < 0.01 and fold change larger than 2 or smaller than -2 were considered as significant. R packages “ChIPseeker” ^57^ was used for peak annotation. Package “ngsplot” ^58^ was used for visualization of cumulated peak signal.

### Statistics

No statistical methods were used to predetermine sample sizes. When comparing two groups, *p* values were determined by two-sided unpaired *t*-test. When comparing more than two groups, *p* values were determined by one-way ANOVA with Tukey’s test. Values are presented as mean ± SEM. The animals were randomly assigned to each experimental or control group. Graphs were generated by the Prism 8.

### Data availability

The source data for the RNA-seq study are available in the NCBI Gene Expression Omnibus (GEO) repository under accession number GSE14659. All other data that support the finding of this study are available from the corresponding author upon reasonable request.

## Acknowledgments

We are grateful to members of the Artandi lab, and to Hosu Sin, Julien Sage, Roel Nusse, Dean Felsher and Margaret Fuller for critical comments. We thank Pauline Chu for expert assistance with histology. Confocal imaging analysis was performed in the Stanford Cell Sciences Imaging Facility. Cell sorting/flow cytometry analysis for this project was performed using the Stanford Shared FACS Facility. Next generation sequencing for this project was performed using the Stanford Functional Genomics Facility. This work was supported by NIH grants CA197563 and AG056575 (to S.E.A) and CA209919 (to H.Y.C.). K.H. was supported by a Japan Society for the Promotion of Science Overseas Research Fellowship.

## Contributions

Conceptualization, K.H. and S.E.A.; Methodology; K.H., Y.Z., R.C., P.C., and S.E.A.; Formal analysis, K.H., Y.Z., A.G., and Y.W.; Investigation, K.H., L.C., R.C., and P.C.; Writing original draft, K.H. and S.E.A.; Supervision, S.E.A., H.Y.C.; Funding acquisition, K.H., S.E.A., and H.Y.C.

## Author information

H.Y.C. is a co-founder of Accent Therapeutics, Boundless Bio, and an advisor to 10x Genomics, Arsenal Biosciences, and Spring Discovery. Correspondence and requests for materials should be addressed to S.E.A.

## Figure legends

**Extended Data Figure 1. TERT expression in spermatogonia subpopulations. (a)** Spermatogenesis: seminiferous tubules are composed of four layers of male germ cells. Undifferentiated and differentiating spermatogonia located in the basement layer undergo mitosis. They translocate to the second layer when entering into meiosis and differentiate into spermatocytes. Haploid spermatids produced by meiosis move toward luminal side during post-meiotic maturation. **(b)** A functional, morphological, and gene expression heterogeneity in spermatogonia subpopulations. **(c)** Flowcytometry analysis of wild-type testicular cells stained with α6-Integrin, MCAM, and KIT. α6-Integrin-high cells were further separated into US-h, US-m, and DS based on KIT and MCAM expression. **(d)** qRT-PCR of *Tert* and spermatogonia markers in FACS-sorted US-h, US-m, and DS. Expression was normalized with *Actb* (n=3 mice per group). Data are represented as Mean ± SEM. **(e)** Flowcytometry analysis of testicular cells from *Tert*^*Tdtomato/+*^ mouse. α6-Integrin-high cells were gated and further separated according to KIT and MCAM expression. *Tert-Tdtomato* expression was compared in US-h, US-m, and DS.

**Extended Data Figure 2. Marking and linage tracing of TERT**^**+**^ **cells. (a)** Immunofluorescence of Tdtomato and E-cadherin, a membrane marker of undifferentiated spermatogonia, in whole-mount seminiferous tubules at day two. Scale bars, 100μm. **(b)** Quantification of Tdtomato^+^ cells in E-cadherin^+^ undifferentiated spermatogonia (n=4 mice per group). Data are represented as Mean ± SEM. **(c)** Whole-mount immunofluorescence of KIT and Tdtomato in a *Tert*^*CreER/+*^*:Rosa*^*lsl-Tdtomato/+*^ testis at two days after high dose tamoxifen injection. Scale bars, 100μm. **(d)** Epifluorescence of Tdtomato in untangled seminiferous tubules at 3-month (3m) or 1-year (1yr) after pulse labeling; injected tamoxifen dose shown (0.25 – 4mg). Scale bars, 5mm. **(e)** Singly isolated Tdtomato^+^ patch. Scale bar, 1mm. **(f)** Cross sections of testes immunostained with antibodies to Tdtomato at 3-month or 1-year after labeling. Scale bar, 50μm.

**Extended Data Figure 3. Conditional deletion of TERT in SSCs using *Tert-flox* mice. (a)** *Tert-flox* targeting strategy and Southern blot strategy. **(b)** Southern blot analysis of genomic DNA from mouse tails. Locations of 5’ and 3’ probes are shown in A. **(c)** TRAP assay using mouse embryonic fibroblasts from *Tert-flox* mice. Cells were transfected with adenovirus to express LacZ or Cre.

**Extended Data Figure 4. (a)** qRT-PCR analysis of *Tert* and *Tertci* in purified Tdtomato^+^ undifferentiated spermatogonia at day 7. Effects of doxycycline addition to drinking water on *Tert* mRNA level. Expression was normalized with *Actb* (n=3 mice per group). **(b)** Telomere repeat amplification protocol (TRAP) assay using purified Tdtomato^+^ undifferentiated spermatogonia from indicated genotypes. **(c)** *Tert*^*CreER/flox*^*:Rosa*^*lsl-Tdtomato/+*^*:Terc*^*-/-*^ mice. **(d)** TRAP assay using whole testes from *Tert*^*CreER/+*^*:Terc*^*+/+*^ and *Tert*^*CreER/+*^*:Terc*^*-/-*^ mouse. **(e)** Epifluorescence of Tdtomato in untangled seminiferous tubules. Scale bars, 5mm. **(f-h)** Quantification of mean patch number (d), mean patch length (e), and total patch length (f) (n=7, 6, 8, 7). Data are represented as Mean ± SEM.

**Extended Data Figure 5. Persistent clone loss phenotype of TERT-deleted spermatogonial stem cells with conditional P53 deletion. (a)** *Tert*^*CreER/flox*^*:Rosa*^*lsl-Ttomato/+*^*:Trp53*^*flox/flox*^ mice. **(b)** *In situ* hybridization against *Trp53* mRNA. Tdtomato^+^ undifferentiated spermatogonia were sorted out at 5 days post labeling and stained. **(c)** Quantification of foci number of *Trp53* mRNA (n=174 and 135 cells from left to right). **(d)** Epifluorescence of Tdtomato in untangled seminiferous tubules. Scale bars, 5mm. **(e-g)** Quantitative data of patch number (e), average patch length (f), and total patch length (g) (n=6, 6, 7,7 mice from left to right). **(h)** Immunofluorescence of γH2AX, PLZF, and Tdtomato in testicular cross sections. *Tert*^*Tdtomato/Tdtomato*^ (G6) mice were used as positive control for γH2AX staining. Scale bars, 30μm. **(i)** Quantification of γH2AX-positive unidiferentiated spermatogonia (n=4 mice per group). Data are represented as Mean ± SEM.

**Extended Data Figure 6. Apoptosis in TERT-deleted undifferentiated spermatogonia. (a)** Immunofluorescence for GFRA1 and TdTomato at 14 days post labeling with tamoxifen. Scale bars, 100μm. **(b)** Epifluorescence for TdTomato in untangled seminiferous tubules at 6 weeks post labeling. Scale bars, 5mm. **(c)** Immunofluorescence for Tdtomato and PLZF, a marker of undifferentiated spermatogonia, using testis cross-sections at 6 weeks after tamoxifen injection. Scale bars, 50μm. **(d)** Immunofluorescence for Tdtomato, PLZF, and BrdU using testis cross sections. Scale bars, 50μm. **(e)** Immunofluorescence of PLZF, Tdtomato, and apoptosis marker, cleaved-PARP (cPARP), on testicular cross-sections at 7 days after labeling. *Tert*^*Tdtomato/Tdtomato*^ (G6) mice were used as positive control for cleaved-PARP staining. Scale bars, 50μm. **(f)** Quantification of Tdtomato^+^ cleaved-PARP^+^ undifferentiated spermatogonia (n=4 mice per group). Data are represented as Mean ± SEM.

**Extended Data Figure 7. Global changes of open chromatin structure in undifferentiated spermatogonia after acute deletion of TERT. (a)** Principal component analysis (PCA) of ATAC-seq data. Colors represent indicated cell types from *Tert*-heterozygous mice. Arrow indicates direction of differentiation. **(b)** Pearson correlation heatmaps of ATAC-seq samples. **(c)** Heatmap representation of 7597 peaks showing significant differences across US-h, US-m, DS, SP, and RS. Each row represents one ATAC-seq peak. Color represents relative ATAC-seq accessibility. **(d and e)** Venn diagrams showing the number of peaks found in spermatogonia subpopulations. **(f)** KEGG pathway analysis of differential peaks between *Tert*^*CreER/+*^ and *Tert*^*CreER/flox*^ US-h.

**Extended Data Figure 8. Changes in open chromatin structure at specific loci after acute deletion of TERT. (a-d)** Read distribution of ATAC-seq data across different cell types. **(a)** *Tert* locus. **(b)** *Ret* and *Gfra1* loci. These genes are dominantly expressed in US-h. **(c)** *Cdh1* and *Zbtb16* loci. These genes are dominantly expressed in both US-h and US-m. **(d)** *Kit* and *Prm1/Prm2* loci. *Kit* is dominantly expressed in DS and *Prm1/Prm2* are dominantly expressed in RS.

**Extended Data Figure 9. Down-regulation of multiple signaling pathways in TERT-deleted US-h. (a)** Volcano plot of RNA-seq data of purified Tdtomato^+^ US-h from *Tert*^*CreER/+*^ and *Tert*^*CreER/flox*^ mice. Genes showing the significant changes (more than 2-fold change and P<0.0001) were colored with red. **(b and c)** gene set enrichment analysis using RNA-seq data of US-h purified from *Tert*^*CreER/+*^ and *Tert*^*CreER/flox*^ mice. nES: normalized enrichment score. **(d)** Immunofluorescence for MYC, PLZF, and Tdtomato at day 7 after tamoxifen labeling. Scale bars, 50μm. **(e)** Quantification of mean signal intensities for PLZF staining in undifferentiated spermatogonia (n=68, 71, 66, 78 cells from left to right). **(f)** qRT-PCR analysis of *Myc* mRNA in purified Tdtomato^+^ US at day 7. Doxycycline was added to drinking water. Expression was normalized with *Actb* (n=3 mice per group). **(g)** qRT-PCR analysis of transgenic human *MYC* mRNA in purified Tdtomato^+^ US at day 7 after tamoxifen labeling. Doxycycline was added to drinking water. Expression was normalized with *Actb* (n=3 mice per group). n.d.; not detected. **(h)** qRT-PCR analysis of MYC pathway genes that were significantly down-regulated in gene set enrichment analysis in figure 4b. Tdtomato^+^ US-h cells were purified at day 7 post labeling by FACS and cDNA was synthesized. Expression level was normalized with *Actb* (n=6,6,5,5,5 mice from left to right).

**Extended Data Figure 10. Model for SSC competition driven by non-canonical functions of TERT**. Summary schematic of catalytic activity-independent functions of TERT in stem cell competition. TERT-deleted SSCs are progressively eliminated from SSC pool through cell competition, reducing the contribution of TERT-deleted SSCs to spermatogenesis over time (left). In wild-type SSCs, TERT promotes competitive clone formation by up-regulating MYC protein. In SSCs lacking TERT, downregulation of MYC protein promotes rapid differentiation, impaired proliferation, and global loss of chromatin accessibility. MYC expression bypasses the requirement for non-canonical TERT, restoring SSC-mediated clone formation.

