## Supplemental figures for "Clonal inactivation of telomerase promotes accelerated stem cell differentiation"

Hasegawa et al. Extended Data Fig. 1

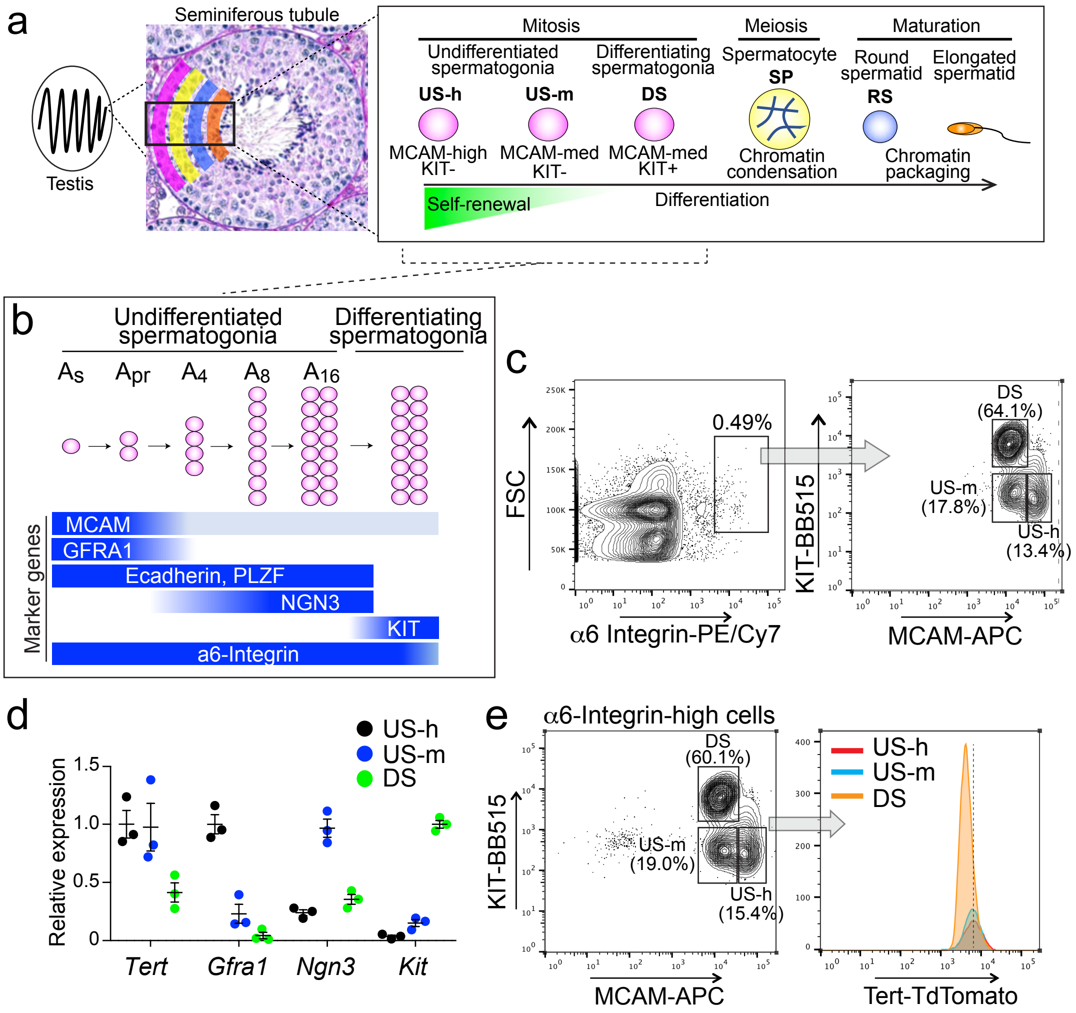

Hasegawa et al. Extended Data Fig. 2

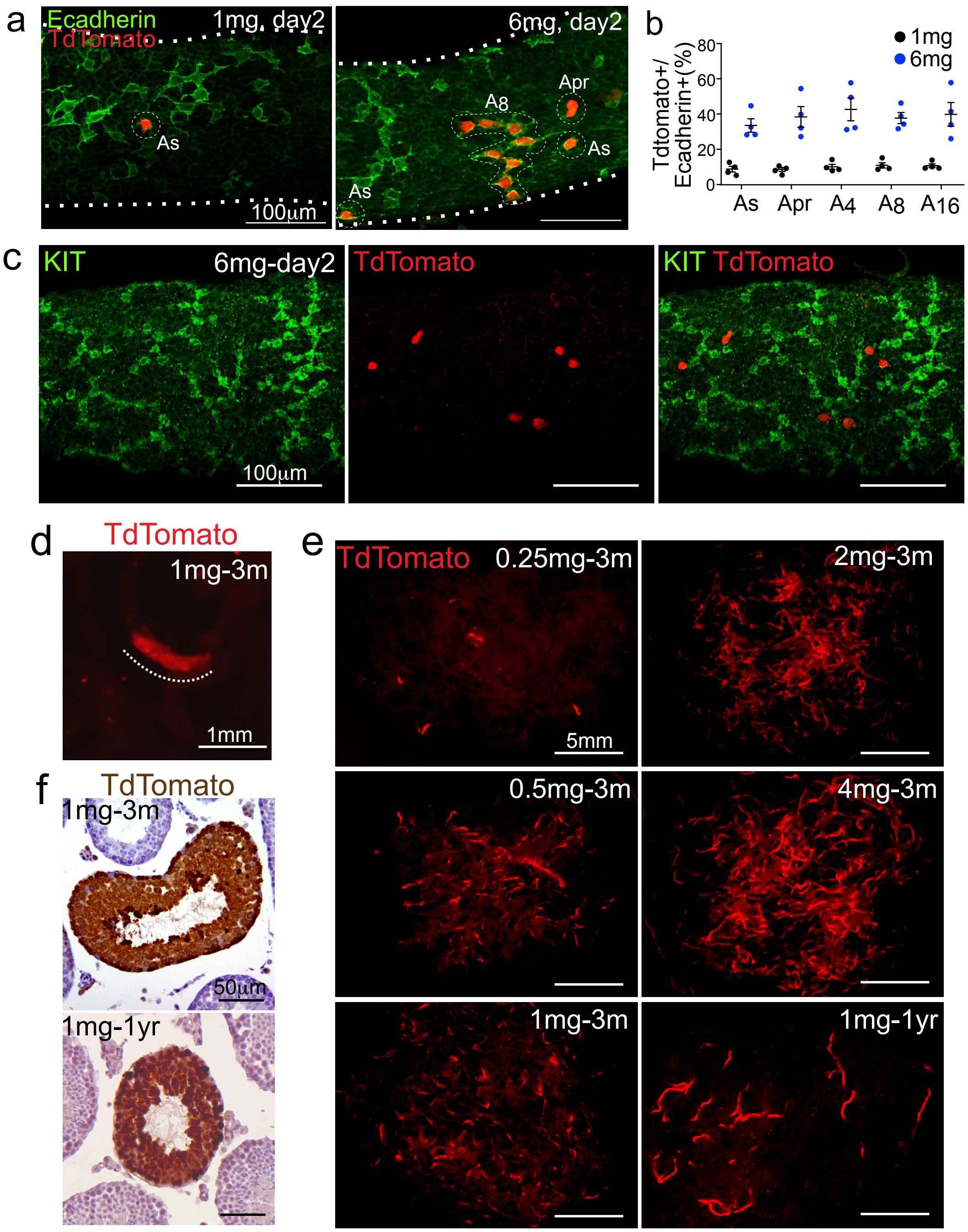

Hasegawa et al. Extended Data Fig. 3

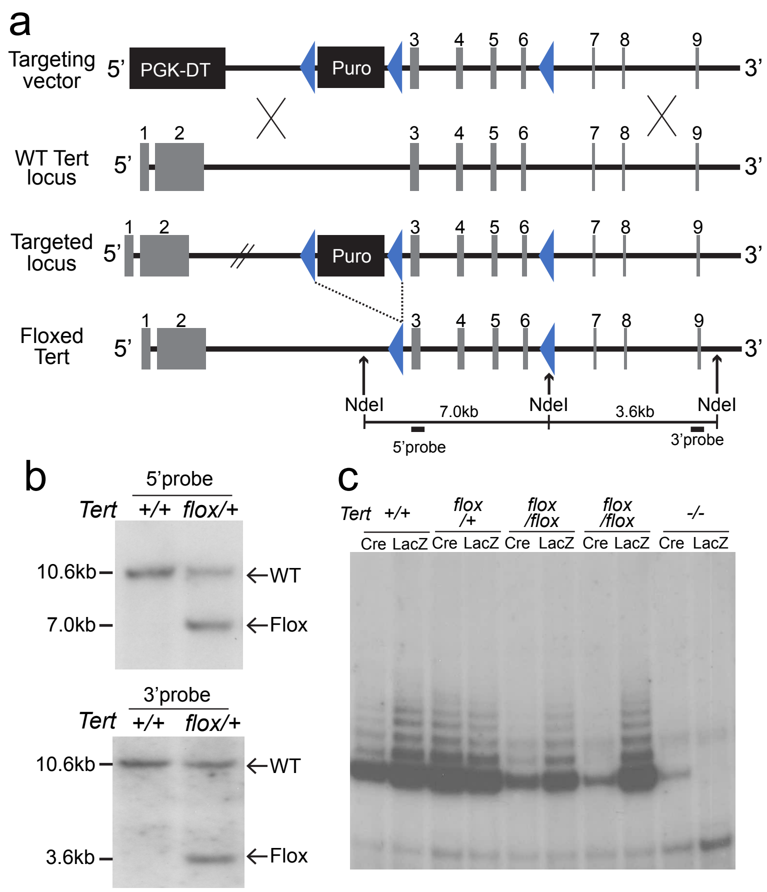

Hasegawa et. al. Extended Data Fig. 4

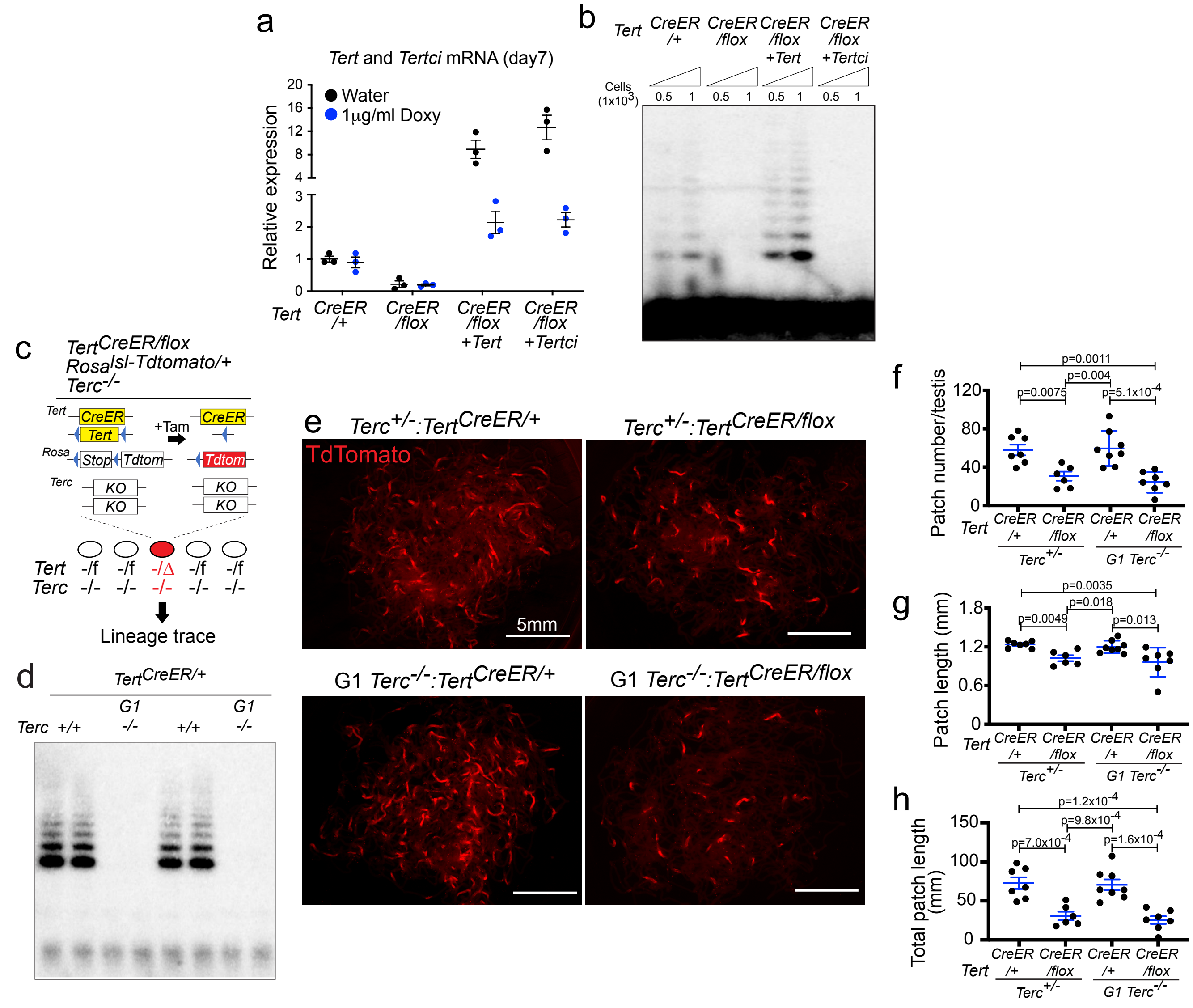

Hasegawa et al.Extended Data Fig. 5

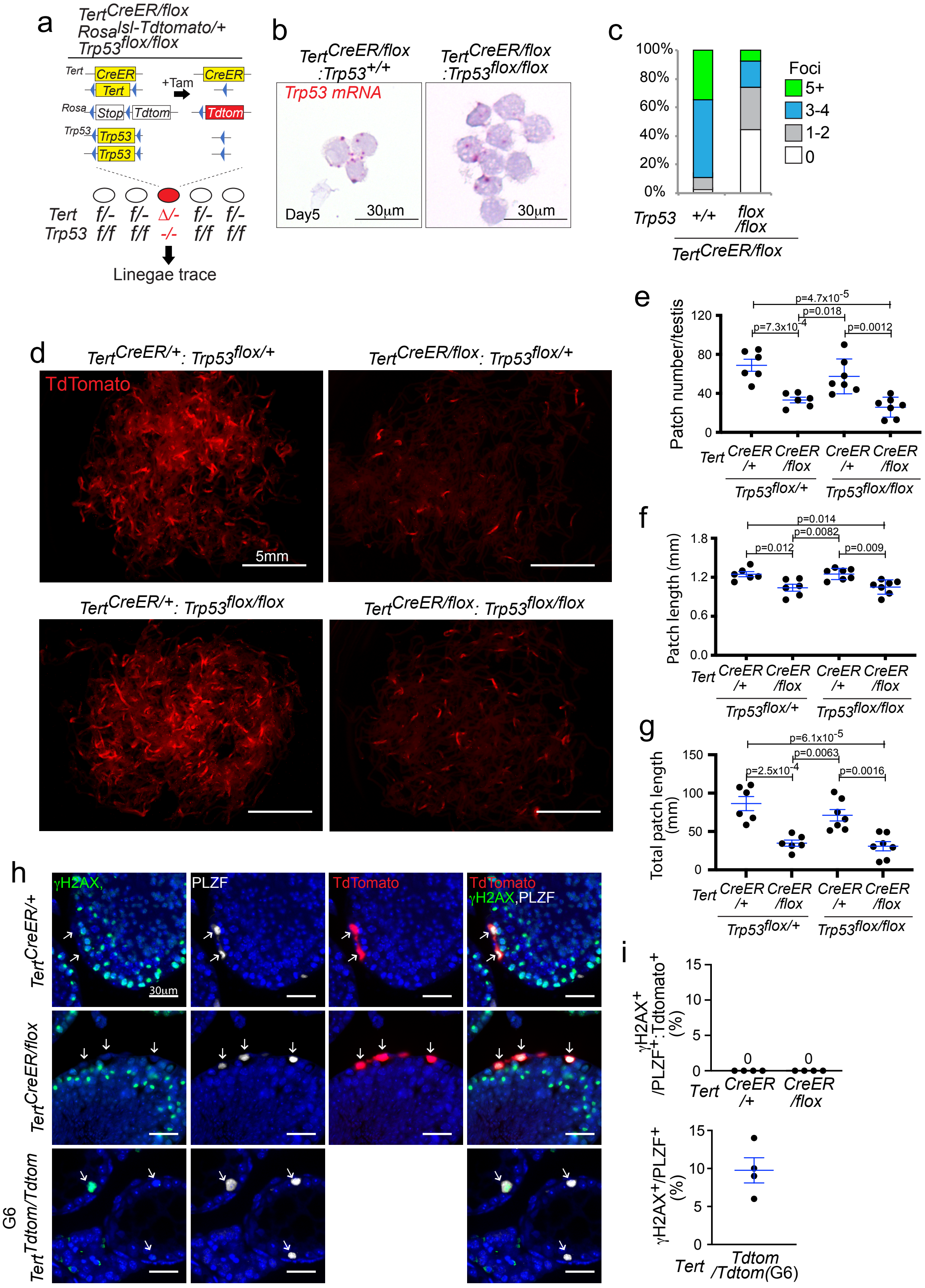

Hasegawa et al. Extended Data Fig. 6

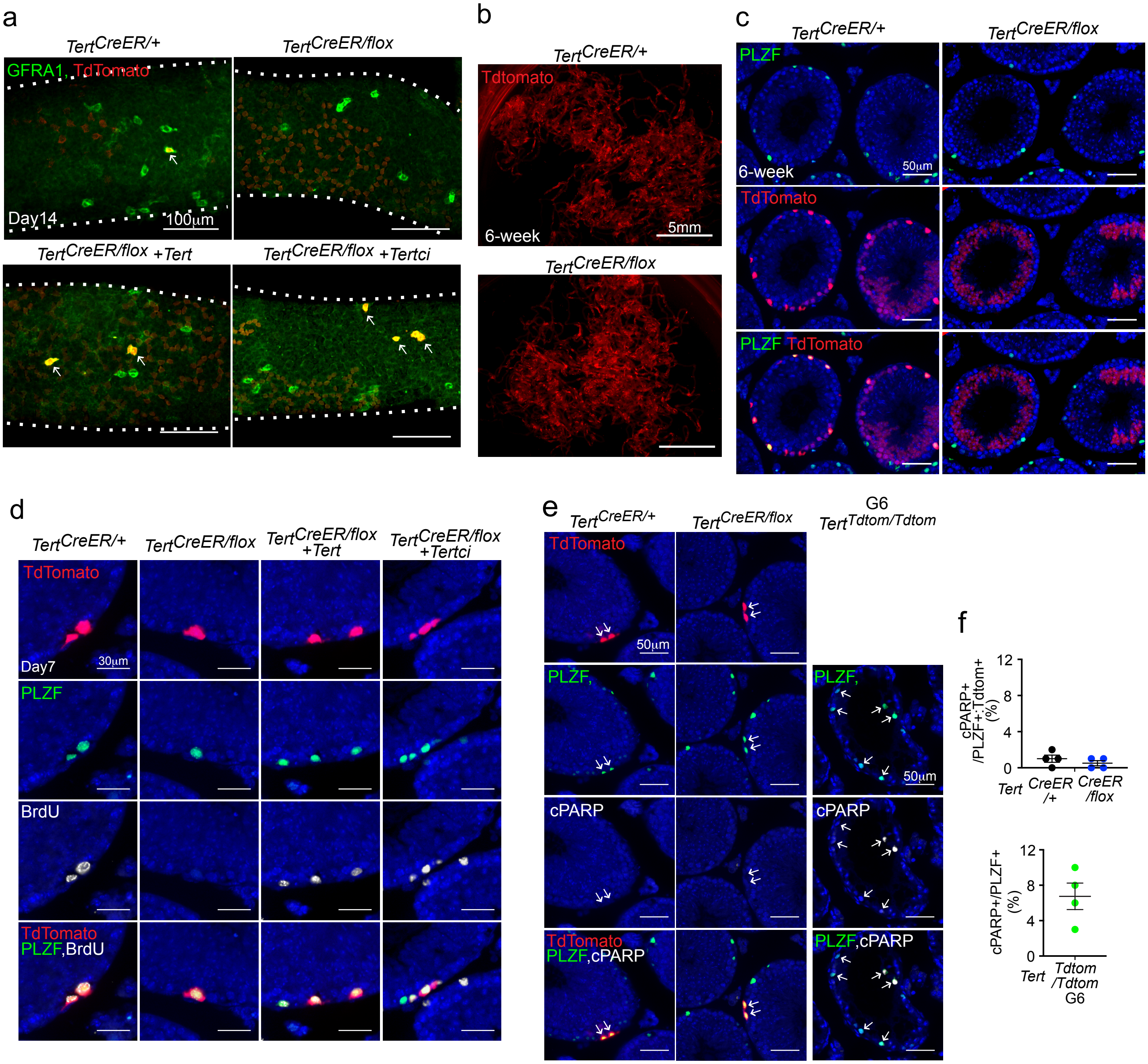

Hasegawa et al. Extended Data Fig.7

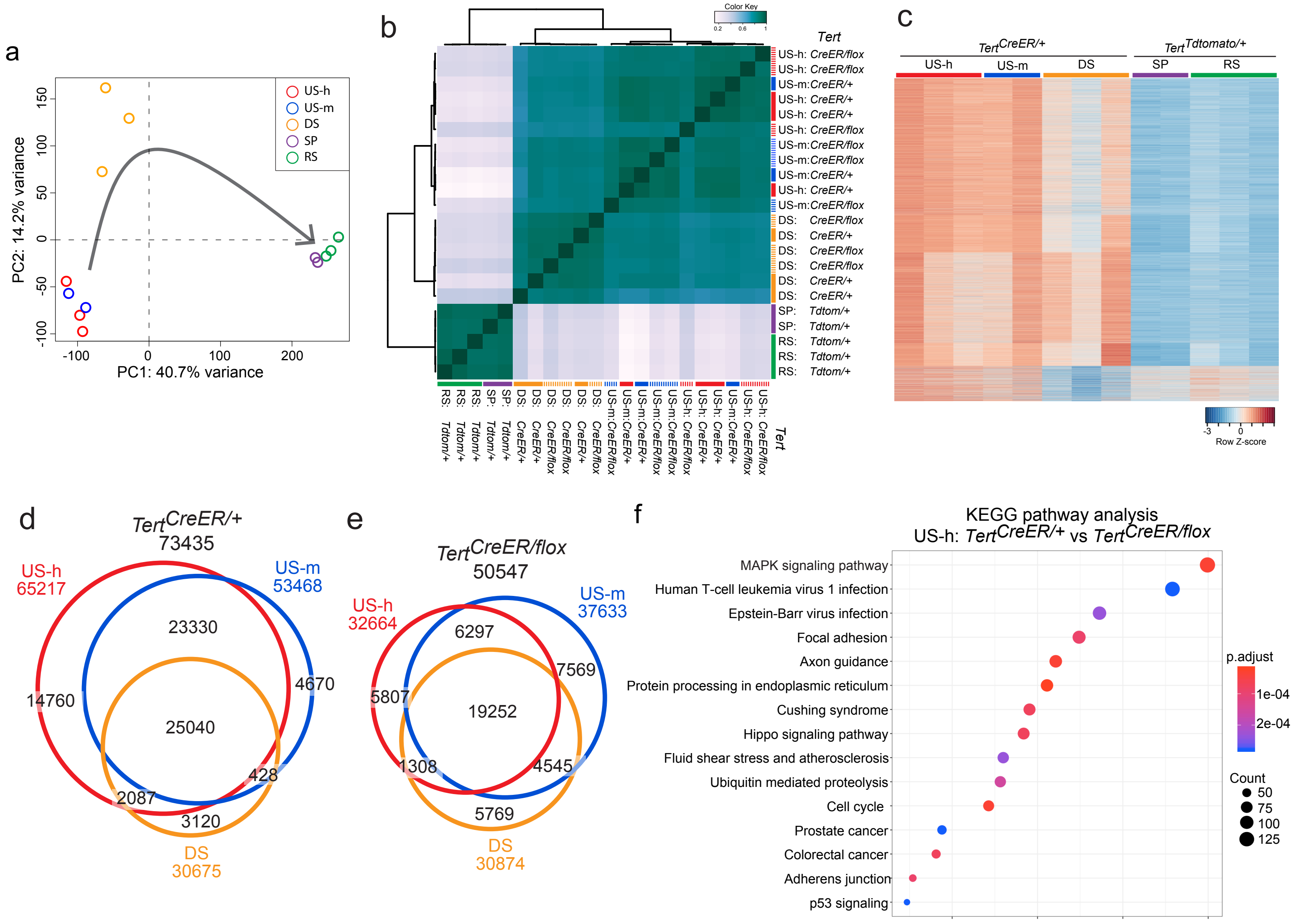

Hasegawa et al. Extended Data Fig. 8

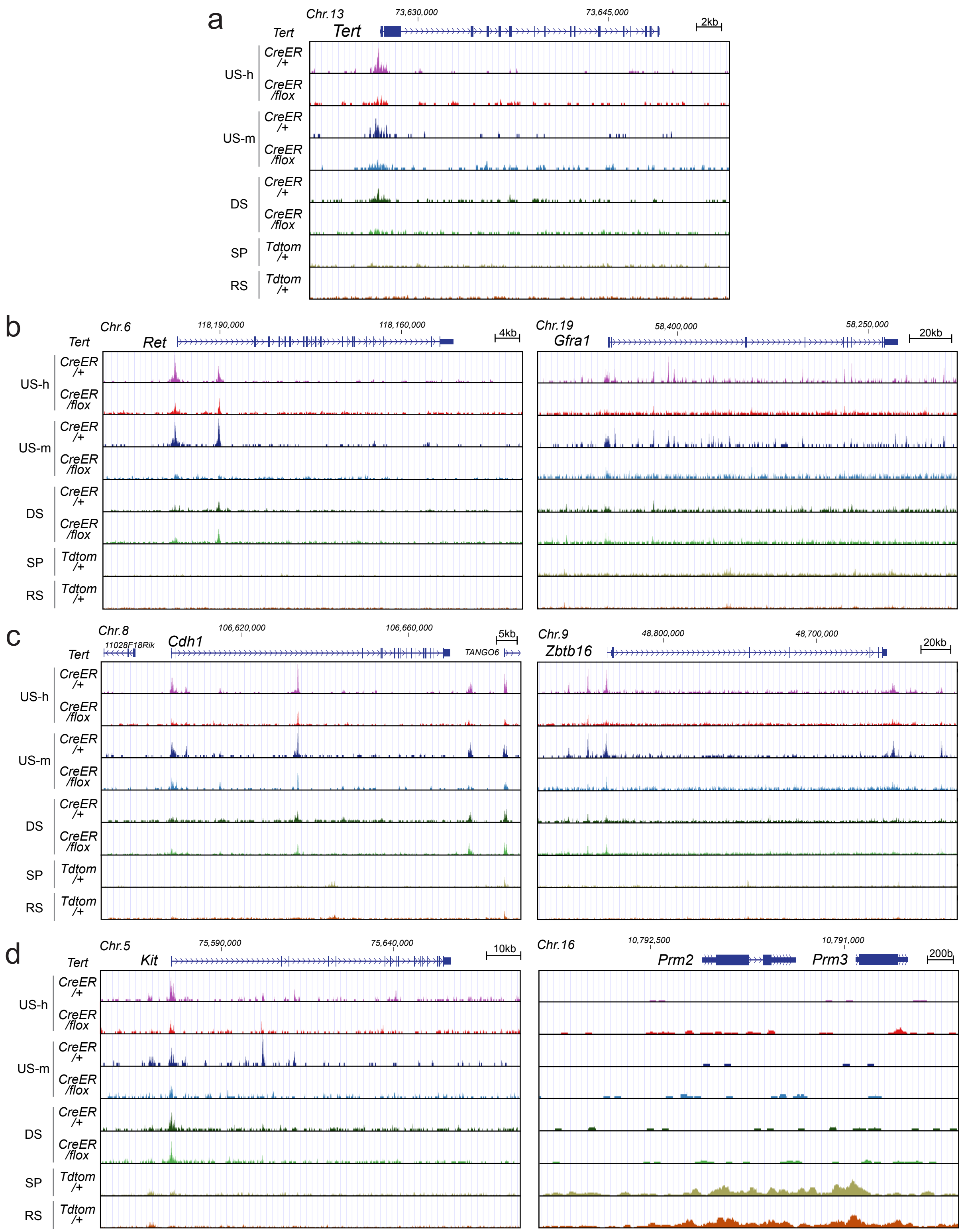

Hasegawa et al. Extended Data Fig.9

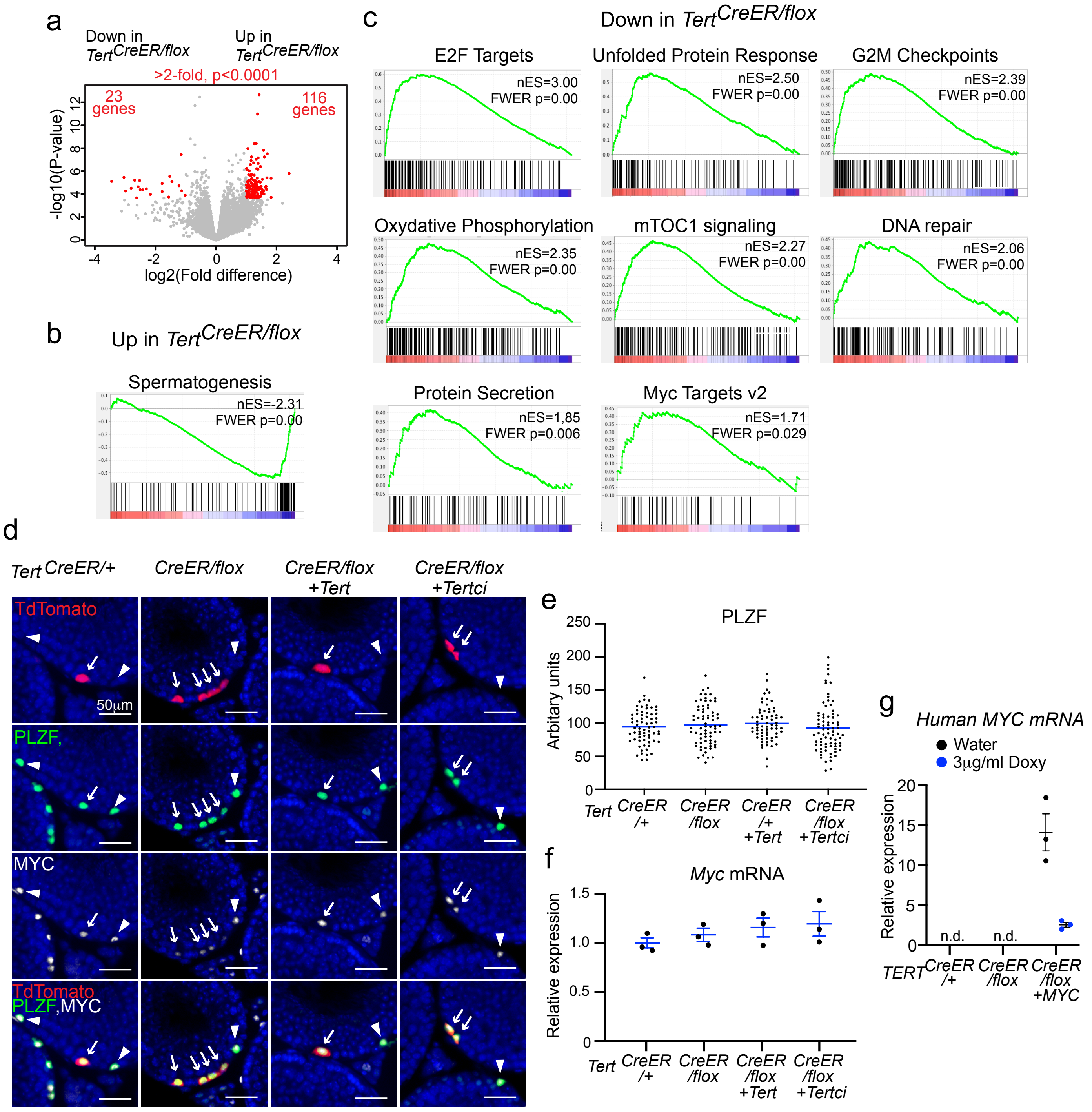

Hasegawa et al. Extended Data Fig.10

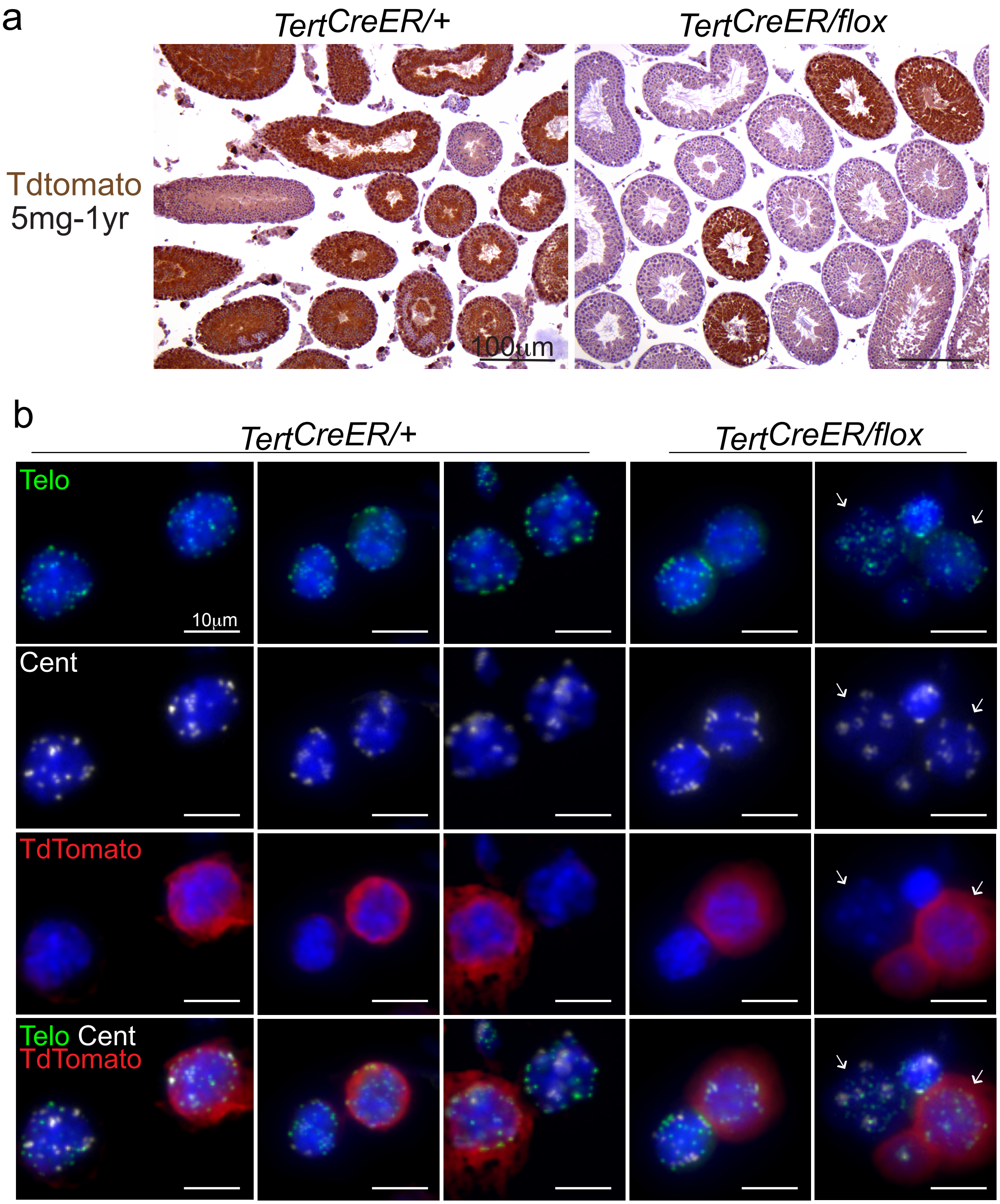
